# Noradrenergic neuromodulation of cholecystokinin interneurons in the basolateral amygdala alters rhythmic activity and restrains fear memory

**DOI:** 10.64898/2025.12.10.693563

**Authors:** Xin Fu, Gajanan Shelkar, Emanuel M. Coleman, Pantelis Antonoudiou, Jamie Maguire, Jeffrey G. Tasker

## Abstract

Norepinephrine (NE) release in the basolateral amygdala (BLA) during emotional arousal plays an essential role in the processing of fear. However, the cell type-specific NE neuromodulation of the fear circuit in the BLA has not been fully resolved. We reported previously a facilitation of fear memory by Gq-coupled receptor induction of repetitive bursting in parvalbumin-expressing interneurons and suppression of gamma oscillations in the BLA. Here, using patch clamp recordings, Cre-dependent DLX-driven intersectional targeting, and genetic manipulations, we demonstrate that NE also activates cholecystokinin (CCK)-expressing interneurons to generate synchronized trains of rhythmic CB1-sensitive IPSCs in BLA principal neurons via Gq-coupled α1A adrenoreceptor activation. The Gq-dependent mechanism in CCK interneurons is generalizable to chemogenetic Gq manipulation and Gq-coupled 5-HT2C serotoninergic receptor activation. We next tested the role of Gq neuromodulation of CCK interneurons in the regulation of BLA network activity associated with the behavioral expression of fear learning by rescued expression of α1A adrenoreceptors or chemogenetic Gq activation selectively in BLA CCK interneurons in a global α1A adrenoreceptor knockout mouse. Restoration of the rhythmic inhibitory synaptic activity via rescue of Gq-coupled receptor signaling in CCK interneurons enhanced LFP theta power in the BLA *in vivo* and decreased fear memory acquisition and recall. These data indicate an inhibitory role for CCK interneuron Gq signaling in fear learning via activation of patterned inhibitory synaptic input to principal neurons and enhanced theta oscillations in the BLA, and reveal a G protein-dependent, receptor-nonselective neuromodulatory mechanism in the BLA that regulates network and behavioral states.

## INTRODUCTION

The basolateral amygdala (BLA) is a cortical-like structure that plays a critical role in the emotional processing of fear (Capogna, 2014; Duvarci & Pare, 2014; Herry & Johansen, 2014; McGaugh, 2004). While plastic changes at glutamatergic synapses on BLA principal neurons have been causally linked to the encoding of associative fear memories (Johansen et al., 2011; Krabbe et al., 2018), accumulating evidence suggests this process is tightly gated by local inhibitory GABAergic circuits (Davis et al., 2017; Krabbe et al., 2018; Krabbe et al., 2019; Trouche et al., 2013; Wolff et al., 2014). GABAergic inhibitory interneurons make up about 20% of the total neuronal population of the BLA and can be classified into three main groups according to a morpho-functional classification, dendritic inhibitory interneurons, perisomatic inhibitory interneurons, and interneuron-selective interneurons (Hájos, 2021; McDonald, 2020; Spampanato et al., 2011). The dendritic inhibitory interneurons comprise mainly the somatostatin-positive interneurons, which project their axons to the distal dendrites of the principal cells to regulate synaptic integration (Muller et al., 2007). The perisomatic interneurons include the cholecystokinin (CCK)-expressing and parvalbumin (PV)-expressing basket cells, which predominantly innervate the soma and the proximal dendrites of principal neurons and control principal neuron spike timing and synchronization (Freund & Katona, 2007; Veres et al., 2017; Woodruff & Sah, 2007), and the PV-expressing chandelier cells that form axo-axonic synapses on the initial segment of principal neurons. Interneuron-selective interneurons, including the vasoactive intestinal peptide (VIP)-positive neurons, specifically target other interneurons to disinhibit principal cells (Krabbe et al., 2019). Differential engagement of distinct inhibitory interneurons in the BLA may serve to control different BLA-dependent behaviors, including fear memory formation.

The BLA and the medial prefrontal cortex (mPFC) are key nodes for emotional processing, including fear learning. Oscillatory states within and between these regions are largely shaped by local inhibitory interneurons and have been shown to govern the behavioral expression of fear (Herry & Johansen, 2014; Karalis et al., 2016; Totty & Maren, 2022). Theta oscillations (4–12 Hz) particularly in BLA-mPFC circuits have been implicated in driving the behavioral expression of fear (Davis et al., 2017; Karalis et al., 2016; Lesting et al., 2013; Lesting et al., 2011; Likhtik et al., 2014; Seidenbecher et al., 2003; Stujenske et al., 2014; Taub et al., 2018). Rhythmic inhibitory postsynaptic currents (IPSCs) with the capacity to entrain theta activity in local field potentials have been shown in the hippocampus and BLA (Bratsch-Prince et al., 2023; Cobb et al., 1995), but the mechanisms underlying theta pacemaker activity in perisomatic interneurons are not well understood. Parvalbumin-expressing (PV) basket cells have been shown to play a critical role in controlling theta and gamma oscillations in the BLA (Antonoudiou et al., 2022) at least partially by generating repetitive bursting activity (Fu et al., 2022), including theta oscillations associated with the behavioral expression of fear (Davis et al., 2017).

The PV and CCK basket cells are differentially regulated by glutamatergic inputs and subcortical neuromodulatory signals (Andrási et al., 2017; Freund & Katona, 2007). CCK, but not PV, basket cell inhibitory neurotransmission is suppressed presynaptically by cannabinoid signaling (Varga et al., 2010; Vogel et al., 2016; Yoshida et al., 2011), the type 1 cannabinoid receptors (CB1Rs) being selectively and abundantly expressed on the axon terminals of the CCK basket cells (Katona et al., 2001; Yoshida et al., 2011). While the role of CCK basket cells in fear regulation has been inferred from studies manipulating CB1 signaling (Atsak et al., 2015; Campolongo et al., 2009; Di et al., 2016; Morena et al., 2016), the specific contribution of neuromodulatory control of CCK cells to the regulation of fear memories remains largely unknown.

Emotional salience facilitates memory formation, a phenomenon largely mediated by noradrenergic signaling in the BLA (McGaugh, 2004). The BLA receives a robust noradrenergic innervation from the locus coeruleus (LC) (Giustino & Maren, 2018), and stressful stimuli, like foot shocks, strongly stimulate the release of NE in the BLA (McIntyre et al., 2002). In addition to converging evidence showing that activation of β adrenoreceptors boosts BLA principal neuron activity (Giustino et al., 2020; McCall et al., 2017) and enhances the consolidation of fear memory (McGaugh, 2004), NE has also been found to facilitate inhibitory synaptic transmission through α1A adrenoreceptors in the BLA of juvenile rats (Braga et al., 2004), which may be due in part to the activation of regular spiking interneurons in the BLA (Kaneko et al., 2008). However, the behavioral role of α1 adrenoreceptors in fear memory formation remains controversial, as studies using the general α1 antagonist prazosin have yielded inconsistent results (Ferry et al., 1999; Gazarini et al., 2013; Lazzaro et al., 2010; Lucas et al., 2019). This is likely due to the expression of α1 adrenoreceptors in both inhibitory neurons and principal neurons as well as a lack of specificity regarding receptor subtype, cell type, and brain region. Thus, it remains unclear how NE modulates specific inhibitory circuits in the BLA to regulate fear memory formation.

We reported previously that GABAergic interneurons, but not principal neurons, in the BLA express the α1A adrenoreceptor subtype and that the activation of Gq-coupled receptors, including α1A adrenoreceptors, stimulates subtypes of inhibitory interneurons that generate two dissociable patterns of inhibitory synaptic responses in the BLA principal neurons. Thus, Gq-coupled receptor activation elicits a low-frequency, phasic pattern of activity in PV basket cells that generates repetitive bursts of high-frequency inhibitory postsynaptic currents (IPSCs) in the BLA principal neurons, depresses gamma-frequency oscillations in the BLA, and facilitates fear memory formation (Fu et al., 2022). In addition, we observed a second stereotyped IPSC response, which we demonstrate here to be mediated by CCK basket cells using a combination of *ex vivo* and *in vivo* electrophysiology, intersectional viral targeting, genetic manipulations, and behavioral analysis. We show that neuromodulation of CCK basket cells modulates theta-band oscillations in the BLA and influences fear learning.

## METHODS

### Animals

Mice were maintained in an AALAC-approved, temperature-controlled animal facility on a 12-h light/dark cycle with food and water provided *ad libitum*. CCK-ires-Cre (Cat. 012706) and adra1A KO mice (Cat. 005039) were purchased from Jackson Laboratories and bred in-house to establish colonies. Heterozygous GAD67-eGFP mice were purchased from Riken BioResource center (Tamamaki et al., 2003) and back-crossed for more than 5 generations with wildtype C57BL/6 mice. All procedures were approved by the Tulane University and Tufts University Institutional Animal Care and Use Committees and were conducted in accordance with Public Health Service guidelines for the use of animals in research.

### AAV virus development

For cloning of Cre-dependent hDlx AAV virus, we amplified the hM3D-mCherry and mCherry coding sequence from the plasmid pAAV-hSyn-DIO-hM3D(Gq)-mCherry (a gift from Bryan Roth, Addgene # 44361) (Krashes et al., 2011) and pAAV-hSyn-DIO-mCherry (also a gift from Bryan Roth, Addgene # 50459) and cloned into pAAV-hDlx-Flex-GFP vector backbone (a gift from Gorden Fishell, Addgene #83895) (Dimidschstein et al., 2016) at AccI and NheI cloning sites. The coding sequence of adra1A was synthesized from Bio Basic Inc. and cloned into a pAAV-hDlx-Flex backbone. AAV virus from Vigene Biosciences Company was further packaged in AAVdj serotype. All hDlx AAV viruses were diluted to the range of 10^11^ to 10^12^ viral genome per ml with virus dilution buffer containing 350 mM NaCl and 5% D-Sorbitol in phosphate-buffered saline (PBS).

### Intracerebral virus injection

*For intracellular recordings:* Four- to 6-week-old male mice were anesthetized by intraperitoneal (I.P.) injection of ketamine/xylazine (100 mg/kg) and placed in a stereotaxic frame (Narishige, SR-6N). The scalp was cut along the midline and the skull was exposed and cleaned. Two burr holes were perforated above the BLA with a Foredom drill (HP4-917). Mice were then injected bilaterally with 350 nl of virus into the BLA (AP: −0.8, ML: 3.05, DV: 4.4) using a 33-gauge Hamilton syringe (10 µL) connected to a micropump (World Precision Instrument, UMP-2) and controller (Micro4) at a flow rate of 100 nL per min. After waiting for 5 min following virus injection to minimize virus spread up the needle track, the injection needle was then slowly retracted from the brain. After surgery, the scalp was sealed with Vetbond, a triple antibiotic ointment was applied, and an analgesic (Buprenorphine, 0.05 mg/kg) was injected I.P.

*For extracellular recordings and drug application:* Eight- to 10-week-old mice were anesthetized with I.P. ketamine/Xylazine (100 mg/kg; 10 mg/kg) and placed in a mouse stereotaxic frame (World Precision Instruments, 502600) over a warm heating pad. Lacri-lube was placed over the subjects’ eyes and slow-release buprenorphine (Buprenorphine SR-LAB, 0.5 mg/kg) was administered subcutaneously for post-operative analgesia. The scalp was shaved, cleaned with ethanol and betadine (3x), then cut along the midline to expose the skull. The skull was leveled and burr holes were manually drilled above the BLA, and 350 nL of virus was injected into the BLA (relative to bregma: AP −1.35, ML ± 3.3, DV −5.1) at a flow rate of 100 nL per minute. After waiting 10 min following injection to minimize viral spread up the needle track, the injection needle was then slowly retracted from the brain. Mice were simultaneously implanted with an EEG/LFP prefabricated headmount (Pinnacle Technology Inc, #8201) customized in-house with a depth electrode which was placed into the BLA (relative to bregma: AP −1.35, ML ± 3.3, DV −5.1) and mounted to the skull using stainless steel screws that acted as ground, reference, and frontal cortex EEG. The headmount was fixed to the skull using dental cement and allowed to cure before removing the animal from the stereotaxic frame. The mice were allowed to recover for a minimum of one week prior to experimentation.

### Ex vivo electrophysiology

*Brain slice preparation:* Coronal brain slices containing the BLA were collected from 6- to 9-week-old male mice for *ex vivo* electrophysiological recordings. The mice were decapitated in a restraining plastic cone (DecapiCone, Braintree Scientific) and their brains were extracted and immersed in ice-cold, oxygenated cutting solution containing the following (in mM): 252 sucrose, 2.5 KCl, 26 NaHCO_3_, 1 CaCl_2_, 5 MgCl_2_, 1.25 NaH_2_PO_4_, 10 glucose (Barsy et al., 2017). The brains were trimmed and glued to the chuck of a Leica VT-1200 vibratome (Leica Microsystems) and 300 µm-thick coronal slices were sectioned. Slices were transferred to a holding chamber filled with oxygenated recording artificial cerebrospinal fluid (aCSF) containing (in mM): 126 NaCl, 2.5 KCl, 1.25 NaH_2_PO_4_, 1.3 MgCl_2_, 2.5 CaCl_2_, 26 NaHCO_3_, and 10 glucose. They were maintained in the holding chamber at 34°C for 30 min before decreasing the chamber temperature to ∼20°C.

*Patch clamp recording:* Slices were bisected down the midline and hemi-slices were transferred one-at-a-time from the holding chamber to a submerged recording chamber mounted on the fixed stage of an Olympus BX51WI fluorescence microscope equipped with differential interference contrast (DIC) illumination. The slices in the recording chamber were continuously perfused at a rate of 2 mL/min with recording aCSF maintained at 32-34°C and continuously aerated with 95% O_2_/5% CO_2_. Whole-cell patch clamp recordings were performed in putative principal neurons in the BLA. Glass pipettes with a resistance of 1.6-2.5 MΩ were pulled from borosilicate glass (ID 1.2mm, OD 1.65mm) on a horizontal puller (Sutter P-97) and filled with an intracellular patch solution containing (in mM): 110 CsCl, 30 potassium gluconate, 1.1 EGTA, 10 HEPES, 0.1 CaCl_2_, 4 Mg-ATP, 0.3 Na-GTP, 4 QX-314; pH was adjusted to 7.25 with CsOH and the solution had a final osmolarity of ∼ 290 mOsm. DNQX, APV, TTX, Prazosin, WB 4101, A61603, CNO, and NE were delivered at the concentrations indicated via the perfusion bath. Slices were pre-incubated in aCSF containing ω-agatoxin (0.5 µM, 30 min), ω-conotoxin (0.5 µM, 30 min), or YM 254890 (10 µM, 20 min) to block P/Q-type calcium channels, N-type calcium channels and Gα_q/11_ activity, respectively (Owen et al., 2013; Takasaki et al., 2004). Series resistance was normally below 10 MΩ immediately after breaking through the membrane and was continuously monitored. Cells were discarded if the series resistance exceeded 20 MΩ.

### In vivo electrophysiology

Local field potential (LFP) recordings were performed in awake, freely behaving wild-type C57BL/6J and CCK-ires-Cre mice using prefabricated headmounts (Pinnacle Technology, #8201) customized with a BLA depth electrode and cannula (Plastics One). BLA LFP recordings were acquired at 4KHz with 100x amplification through an insulated LFP depth electrode implanted in the ipsilateral BLA. All mice were allowed to habituate to the recording chamber for at least 1 h prior to recording. In CCK-ires-Cre animals expressing hM3D in BLA CCK interneurons, baseline and treatment conditions were recorded for 60 min each. Treatments consisted of 0.5 μL infusion of saline or CNO (1 mM in isotonic saline) into the BLA at a rate of 0.1 μL per min. Analysis was performed using custom in-house scripts (SAKE, GitHub). The vehicle and CNO responses were normalized to baseline and the data represented as CNO responses relative to vehicle.

### Fear conditioning

Three weeks after virus injection, mice were single housed and handled for ≥ five days before undergoing the fear conditioning paradigm with the Video Fear Conditioning System in a sound attenuated chamber (MED Associates, Inc.). Each chamber is equipped with a metal stainless-steel grid connected to a shock generator (ENV414S Aversive Stimulator). The fear conditioning paradigm consisted of 7 exposures to a continuous tone (7 kHz, 80 db, 30 s duration) as the conditioning stimulus (CS), each of which was co-terminated with the unconditioned aversive stimulus (US) consisting of an electric foot shock (0.7 mA, 2 s duration). The CS-US stimuli were presented at a randomized intertrial interval (ITI, 30-180 s, average = 110 s) in one context, context A. Twenty-four hours later, on day 2, mice were tested for fear retrieval in a different context, context B, with a planar floor and a black plastic hinged A-frame insert. During fear memory retrieval, five presentations of CS alone were delivered with an inter-stimulus interval of 60 s. Behavior was recorded with an infrared camera and analyzed with Video Freeze software (Med Associates, Inc.). Mice were considered to exhibit freezing behavior if no movement other than respiration was detected for ≥ 2 s. Chambers were cleaned with either 70% ethanol or 3% acetic acid before each session of fear conditioning or fear memory retrieval.

### Histology

#### Perfusion and cryosectioning

Two weeks after AAV virus injection, adra1A KO, CCK-ires-Cre, CCK-ires-Cre::GAD67-eGFP, and CCK-ires-Cre::adra1A KO mice were deeply anesthetized with ketamine/xylazine (300 mg/kg) and perfused transcardially with 10 mL of ice-cold PBS (pH 7.4) followed by 20 mL of 4% paraformaldehyde (PFA) in PBS. Brains were dissected out, postfixed for 3 h in 4% PFA in PBS, and cryopreserved with 30% sucrose in PBS for 24 h at 4°C, or until the brain sunk to the bottom of the container. Frozen coronal sections (45 µm) were cut on a cryostat (Leica) and harvested in 24-well plates filled with PBS at room temperature.

#### Confocal imaging

Sections from virus-injected CCK-ires-Cre, CCK-ires-Cre::GAD67-eGFP, and CCK-ires-Cre::adra1A KO mice containing the BLA were selected, rinsed with PBS (3 x 5 min), and mounted on gel-coated slides. Confocal images were acquired with a Nikon A1 confocal microscope to capture the DAPI (excitation 405 nm, emission 450 nm), GFP (excitation 488 nm, emission 525 nm), and mCherry (excitation 561 nm, emission 595 nm) signals. For the analysis of colocalization, z-stack pictures were imaged under a 40x oil-immersion objective with a step increment of 1.5 µm. The number of BLA cells containing colocalized markers were first quantified from merged maximal intensity images from different channels, and then confirmed in z-stack images with ImageJ (NIH).

#### X-gal staining

Sections from adra1A KO mice were rinsed with PBS (3 x 5 min) and incubated in a β-gal staining solution (Roche, Ref # 11828673001) overnight. After β-gal staining, sections were then rinsed in PBS (3 x 5 min), mounted on gel-coated glass slides, coverslipped with Permount mounting medium (Fisher Scientific), and allowed to air dry. Bright-field imaging was performed in a Zeiss Axio Scanner and processed and analyzed with ImageJ (NIH). For fluorescence confocal imaging, brain sections were rinsed with PBS (3 x 5 min), mounted on gel coated coverslips, and then imaged first on the confocal microscope (Nikon A1) before incubating them in the β-gal staining solution. After staining with X-gal, the slices were rinsed with PBS (3 x 5 min) and the same regions previously imaged using fluorescence confocal imaging were re-imaged for the X-gal signal with excitation and emission wavelengths of 638 and 700 nm (Levitsky et al., 2013), respectively. The images were then processed and quantified with ImageJ software to determine the ratio of β-gal-positive cells to fluorescent cells with the same procedure described above.

### Data analysis

#### Intracellular analysis

Data were acquired using a Multiclamp 700B amplifier, a Digidata 1440A analog/digital interface, and pClamp 10 software (Molecular Devices). Recordings were filtered at 2 KHz for IPSC recordings and sampled at 50 KHz. Data were analyzed with MiniAnalysis software (SynaptoSoft, NJ) and Clampfit 10 (Molecular Devices). Statistical comparisons were conducted with a paired or unpaired Students’s *t* test or with a one- or two-way ANOVA followed by a *post hoc* Dunnett’s test as appropriate (p < 0.05 with a two-tailed analysis was considered significant).

#### In vivo LFP analysis

LFP data were analyzed as previously described (Antonoudiou et al., 2022; Fu et al., 2022). Signals were band-pass filtered (1-300 Hz, Chebyshev Type II filter), and spectral analysis was performed in MATLAB using publicly accessible custom-made scripts using the fast Fourier transform (Frigo and Johnson 2005). Briefly, recordings were separated into 5-s bins with 50% overlapping segments. The power spectral density for positive frequencies was obtained by applying a Hann window to eliminate spectral leakage. The mains noise (58-60 Hz band) was removed from each bin and replaced using the PCHIP method. Values 5 times larger or smaller than the median were considered outliers and replaced with the nearest bin. The vehicle and CNO responses were normalized to the baseline activity. The data were then imported into GraphPad Prism for statistical analysis. Statistical comparisons at each oscillatory frequency were conducted with one-sample *t* tests (p < 0.05 with a two-tailed analysis was considered significant).

## RESULTS

### NE stimulates CCK basket cells to generate synchronized rhythmic inhibitory postsynaptic currents in BLA principal neurons

In whole-cell recordings from BLA principal neurons, norepinephrine (NE) elicited two distinct bursting patterns of IPSCs mediated by different subpopulations of local inhibitory interneurons, an initial burst of low-frequency IPSCs followed by a repetitive train of bursts of high-frequency IPSCs (**Suppl. Fig. 1**). The repetitive train of IPSC bursts is generated by alpha-1A adrenoreceptor (ARα1A) activation of action potential burst firing in presynaptic PV interneurons, which we described in detail in a previous report (Fu et al., 2022). Briefly, the PV neuron-mediated IPSC bursting response to NE was blocked by the selective P/Q-type calcium channel blocker ω-agatoxin (0.5 µM), which isolated the second type of IPSC burst (**Suppl. Fig. 1**). Here, we focused on the initial NE-stimulated burst of IPSCs in the BLA principal neurons. After blocking glutamatergic transmission with DNQX (20 µM) and APV (40 µM) and the PV neuron-mediated inhibitory input with ω-agatoxin (0.5 µM), NE (100 µM) induced 1-5 inactivating ‘bursts’ of IPSCs in the principal neurons that were characterized by a slow, pacemaker-like rhythmic instantaneous intra-burst frequency of ∼ 4 Hz (range: 2.02 to 5.23 Hz, mean: 3.58 Hz) and a duration of ∼ 70 s (range: 25 to 198.4 s, mean: 69.25 s) (**Fig. 1A-D**). The NE-induced, low-frequency IPSC bursts terminated prior to washout of the NE, suggesting that they desensitized to the NE with time. When multiple IPSC bursts were recorded in a principal neuron, the bursts differed in duration and IPSC amplitudes and frequencies, but showed a similar intra-burst rhythmic IPSC pattern, suggesting that each burst was generated by a different presynaptic GABA interneuron from the same interneuron subpopulation (**Fig. 1C**). The bursts of low-frequency IPSCs were inhibited by blocking spiking activity with tetrodotoxin (0.5 µM) and with the alpha 1A adrenoreceptor antagonist WB4101 (1 µM), and were mimicked by the alpha 1A adrenoreceptor agonist A61603 (2 µM) (**Fig. 1E**). They were also blocked by pre-incubation of the slices in the selective Gα_q_ inhibitor YM-254890 (10 µM) (**Fig. 1E**). Taken together, these data suggest that NE activates α1A adrenoreceptors to stimulate a subtype of BLA local inhibitory interneuron, which in turn generates a single, inactivating burst of rhythmic, low-frequency IPSCs in BLA principal neurons.

**Figure 1.**
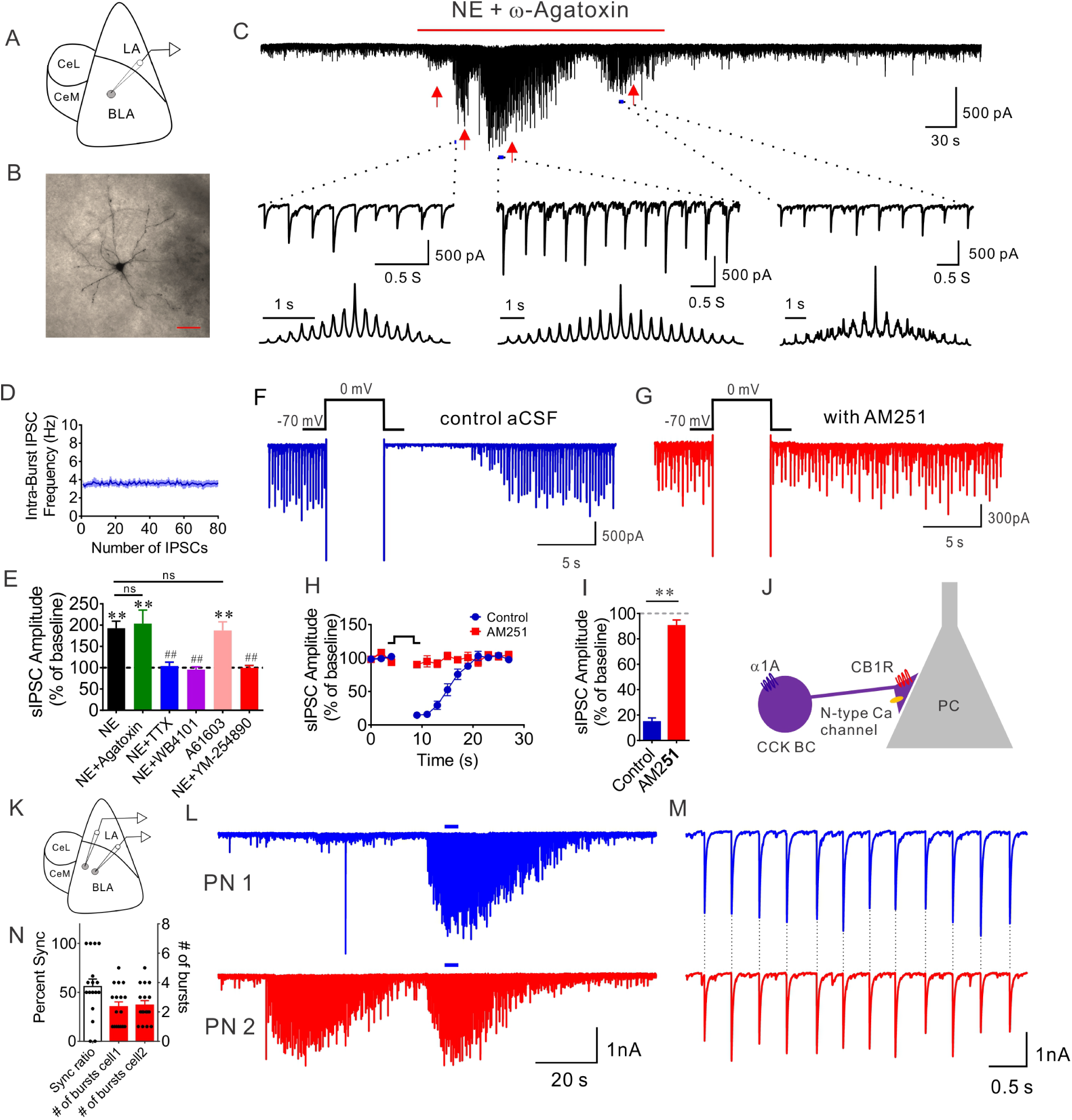
NE stimulates synchronized bursts of low-frequency, rhythmic IPSCs in BLA principal neurons by Gq-coupled alpha1A receptor activation of local CCK interneurons. **A.** Schematic diagram of *ex vivo* BLA recordings. **B**. Biocytin staining of a recorded BLA principal neuron. **C**. A representative recording showing multiple trains of rhythmic IPSCs induced by NE in the presence of the P/Q-type calcium channel blocker ω-agatoxin. Red arrows indicate different IPSC bursts, three of which were expanded below to show the regularity of the individual IPSCs within each burst, with the corresponding autocorrelograms beneath each expanded burst (note different time scales). **D**. Population mean (± SEM) instantaneous frequency of NE-induced IPSCs throughout the course of the bursts (n = 11 bursts from 8 cells). **E**. Mean change in amplitude of NE-induced IPSCs. The NE-induced increase of IPSC amplitude (16 cells from 5 mice) was not affected by blocking P/Q-type calcium channels with ω-agatoxin (12 cells from 5 mice) but was eliminated by blocking neural activity with the sodium channel blocker TTX (10 cells from 5 mice). The NE-induced rhythmic IPSCs were blocked by the α1A adrenoreceptor antagonist WB4101 (7 cells from 4 mice) and mimicked by the α1A adrenoreceptor agonist A61603 (10 cells from 4 mice). Inhibition of Gq activation with YM-254890 also blocked the NE-induced increase in IPSC amplitude (8 cells from 3 mice). Paired Student’s *t* tests: NE vs. baseline, p<0.0001; NE + Agatoxin vs. baseline, p = 0.0071; A61603 vs. baseline, p = 0.0024; ** p < 0.01; one-way ANOVA, F (5, 57) = 6.529, p < 0.0001; Dunnett’s multiple comparisons test: NE vs. NE+Agatoxin, p = 0.99; NE vs. NE+TTX, p = 0.0057; NE vs. NE+WB4101, p = 0.0072; NE vs. A61603, p = 0.99; NE vs. NE+YM-254890, p = 0.007. Values at the peak of IPSC amplitude (time = 7 min in all the treatments) were used to compare ω-agatoxin-insensitive IPSCs. ## p < 0.01. **F, G**. Representative traces showing ω-agatoxin-isolated, NE-induced IPSCs that underwent transient and complete depolarization-induced suppression of inhibition (DSI) (F), which was reversed by *the* CB1 receptor antagonist AM251 (G). **H, I**. Time course (H) and mean change in the IPSC amplitude (I) during DSI of NE-induced IPSCs before (Control, 8 cells from 3 mice) and after CB1 receptor blockade (AM251, 3 cells from 2 mice). Unpaired Student’s *t* test: control vs. AM251, p < 0.0001, ** p < 0.01. **J**. Proposed model of NE modulation of CB1 receptor- and N channel-sensitive CCK basket cell (BC) input to BLA principal cells (PC). **K**. Schematic diagram of paired recordings of BLA principal neurons. **L**. Representative traces showing NE-induced IPSC bursts in a pair of BLA principal neurons recorded in the P/Q calcium channel blocker ω-agatoxin. **M**. Segments of the synchronized bursts of IPSCs (blue bars in M) were expanded to show synchronization of individual IPSCs (vertical dashed lines) between the two principal neurons. **N**. The ratio of synchronization and total number of different subtypes of bursts (mean ± SEM) induced in pairs of principal cells by NE activation of CCK interneurons (17 principal neuron pairs from 6 mice).

The individual IPSCs in the bursts displayed a fast rise time (1.40 ms ± 0.05 ms) and fast 10-90% decay time (18.43 ms ± 1.16 ms, n=11 bursts in 8 cells from 5 mice), which is characteristic of perisomatic inhibitory signals from basket cells in the BLA (Barsy et al., 2017; Wilson et al., 2001), comprised of CCK- and PV-expressing interneurons. Consistent with our previous findings (Fu et al., 2022), the NE-induced IPSC bursts were blocked by the N-type calcium channel antagonist ω-conotoxin (1 µM), but not by the P/Q calcium channel antagonist ω-agatoxin (0.5 µM) (**Fig. 1E**). They were also completely blocked by CB1 receptor activation with WIN 55,212-2 (1 µM) (**Suppl. Fig. 1**) and by endocannabinoid-dependent depolarization-induced suppression of inhibition (DSI), which was reversed by the CB1 receptor antagonist AM251 (10 µM) (**Fig. 1F-I**). Since the CCK basket cells use N-type, but not P/Q-type, calcium channels for GABA release and because CB1 receptors are selectively expressed in axons of CCK, but not PV, basket cells (Freund & Katona, 2007; Katona et al., 2001; Wilson et al., 2001; Yoshida et al., 2011), these data indicated that the NE-induced generation of bursts of low-frequency, rhythmic IPSCs in BLA principal neurons is mediated by the activation of local presynaptic CCK basket cells (**Fig. 1J**).

Since perisomatic basket cells innervate hundreds of postsynaptic principal neurons through their extensive axonal arborizations (Vereczki et al., 2016), we next asked whether the NE-induced, CCK neuron-generated IPSC bursts are synchronized among BLA principal neurons. To address this question, we conducted paired recordings in BLA principal neurons within ∼50 µm of each other and monitored the response to NE of each cell in the pair simultaneously after blocking PV interneuron inputs with ω-agatoxin (**Fig. 1K**). NE application induced one or more synchronized IPSC bursts in both cells in most of the paired recordings, although all bursts seen in both cells of the pair were not always synchronized (**Fig. 1L**). Within the NE-induced synchronized bursts between two principal cells, individual IPSCs in the bursts were also synchronized (**Fig. 1M**). Dividing the total number of CCK-mediated IPSC bursts by the number of synchronized bursts, we calculated that one BLA principal neuron receives, on average, bursting IPSC inputs from 2.4 presumably different presynaptic CCK interneurons (**Fig. 1N**), and that 55.9% of all the IPSC bursts are synchronized between the pairs of recorded cells (17 recorded pairs from 6 mice). These data suggest that NE may regulate synchronous neural output from the BLA by activating perisomatic basket cells to generate synchronous bursts of inhibitory synaptic inputs.

### Expression of α1A adrenoreceptors in CCK interneurons of the BLA

To test whether CCK interneurons express α1A adrenoreceptors as our pharmacological experiments suggested, we first labeled the CCK interneurons by injection of intersectional hDLX virus (AAVdj-hDLX-DIO-mCherry) in the BLA of a CCK-ires-Cre-expressing mouse line (**Fig. 2A**). Consistent with previous studies (Dimidschstein et al., 2016; Liu et al., 2020), injection of a Cre-dependent hDLX virus selectively labeled CCK interneurons in the BLA, since 98.2 ± 1.5% of the mCherry-labeled cells were colocalized with the interneuron marker GAD-GFP (**Fig. 2B, C**).

**Figure 2.**
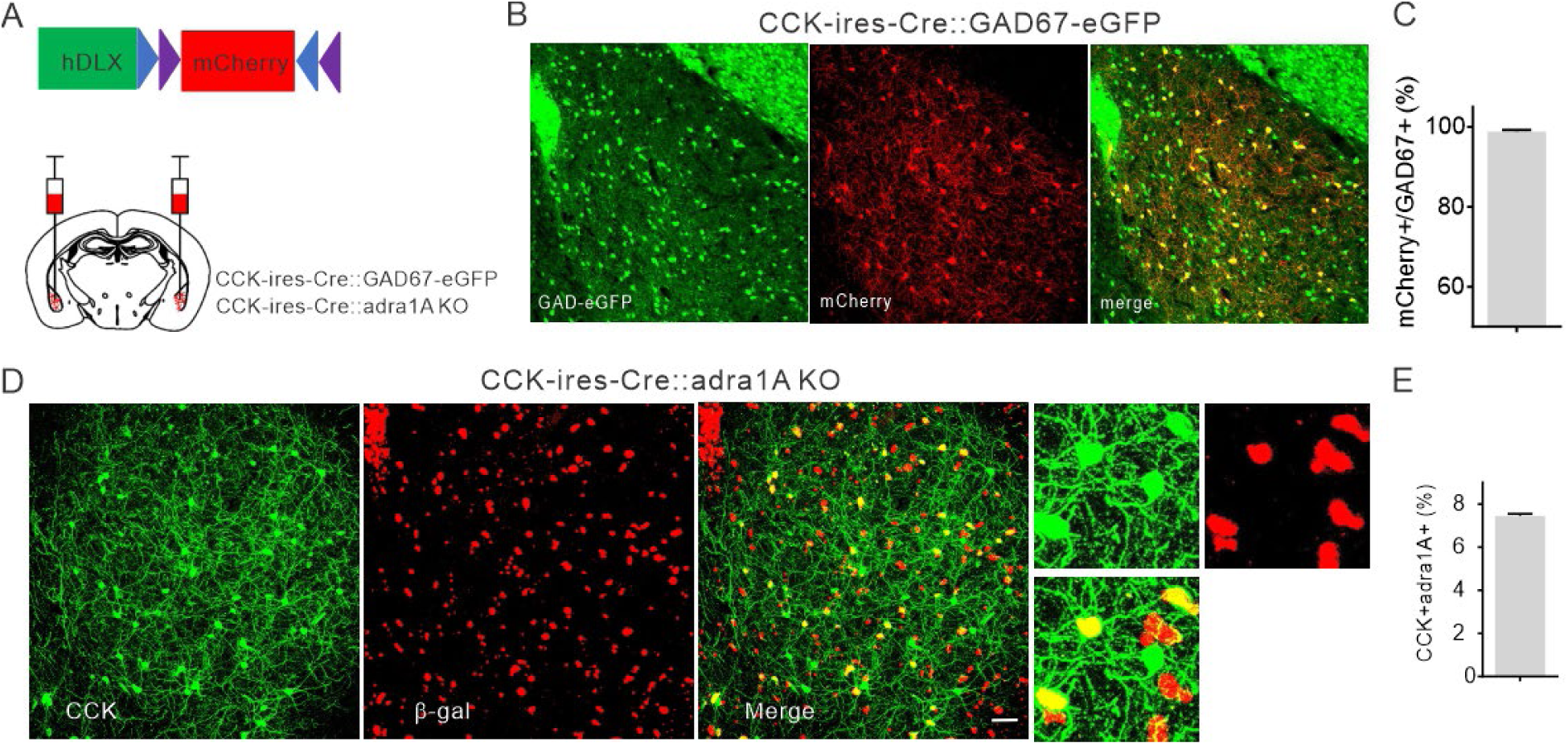
Expression of α1A adrenoreceptors in BLA CCK interneurons. **A**. Schematic diagram showing bilateral injection of Cre-dependent intersectional hDLX-DIO-mCherry virus into the BLA of CCK-ires-Cre::GAD67-eGFP and CCK-ires-Cre::adra1AKO mice. **B**. Representative fluorescence images showing co-expression of mCherry with Gad67-eGFP, which labels nearly all GABAergic interneurons in the BLA. **C**. Quantification of the colocalization of CCK-mCherry and Gad67-eGFP (98.2 ± 1.5%, n=430 cells). **D**. Representative images showing that most of the CCK interneurons labeled with mCherry co-expressed β galactosidase. **E**. Quantification of the colocalization of CCK-mCherry and β-gal (74.5 ± 1.99%, n= 145 cells).

We next examined whether α1A adrenoreceptors are expressed in BLA CCK interneurons. Because the commercially available antibodies against the α1 adrenoreceptors are non-specific (Jensen et al., 2008), we used a global *adra1A* knockout mouse line in which a lacZ gene cassette is knocked into the α1A adrenoreceptor gene locus (adra1AKO) (Rokosh & Simpson, 2002), which allowed us to monitor α1A adrenoreceptor expression using β-galactosidase staining. To co-label CCK interneurons and α1A adrenoreceptor-expressing neurons, we injected the hDLX-DIO-mCherry virus bilaterally into the BLA of a cross between the CCK-ires-Cre mouse and the adra1AKO mouse (CCK-ires-Cre::adra1AKO). After two weeks, we stained for β-galactosidase with X-gal to reveal the cells that express the α1A adrenoreceptors (**Fig. 2D**). A majority of the labeled CCK interneurons (74.5%, **Fig. 2E**) were positive for X-gal. This provides histological evidence for α1A adrenoreceptor expression in CCK basket cells, which supports our data suggesting that the generation of rhythmic IPSC bursts in principal neurons is mediated by α1A adrenoreceptor activation in local CCK interneurons.

### Gq activation of CCK interneurons drives bursts of rhythmic IPSCs in principal cells

Having established the role of CCK interneurons in the generation of the NE-induced bursts of low-frequency IPSCs through activation of Gq-coupled α1A adrenoreceptors, we next tested whether selective chemogenetic Gq activation in CCK interneurons using a designer-receptor (DREADD) strategy (Roth, 2016) would be sufficient to stimulate the same IPSC bursting pattern in the BLA principal neurons. We expressed a Gq-coupled DREADD (hM3D) in the CCK interneurons by bilaterally injecting in the BLA of CCK-ires-Cre-expressing mice an AAV virus that expresses Cre-dependent hM3D under the control of the GABA neuron-specific hDLX promoter (AAVdj-hDLX-DIO-hM3D-mCherry) (Dimidschstein et al., 2016; Liu et al., 2020) (**Fig. 3A**). Two weeks after the intracerebral virus injection, we observed that the expression of Gq-mCherry was also restricted to the interneurons of the BLA (**Fig. 3B, C**). Notably, whole-cell voltage clamp recordings in *ex vivo* slices containing the BLA from virus-injected animals showed that selective activation of Gq-DREADDs in the CCK interneurons with clozapine-N oxide (CNO, 5 µM) predominantly induced bursts of rhythmic, low-frequency IPSCs (2.74 to 7.37 Hz, mean = 4.01 Hz, Fig. 24D-G) in BLA principal neurons (**Fig. 3D-F**) that were very similar to the α1A adrenoreceptor-stimulated IPSC bursts induced by NE (see Fig. 1). In a subset of recorded neurons (2 out of 10 cells), we also observed non-inactivating repetitive bursts of IPSCs with an accelerating intra-burst frequency in addition to the CCK interneuron-mediated rhythmic IPSCs (**Suppl. Fig. 2A**). These bursts were likely to be mediated by Gq activation in PV interneurons (Fu et al., 2022) and are consistent with a previous report showing that intersectional targeting of CCK interneurons with CCK-ires-Cre can label a diverse population of interneurons, including the PV-positive interneurons in the BLA (Rovira-Esteban et al., 2019). Consistent with the double dissociation of two NE-induced patterns of IPSCs with calcium channel blockers, the rhythmic IPSC bursts generated by hM3D activation in CCK interneurons were abolished by blocking N-type calcium channels with ω-conotoxin (1 µM) and by blocking Gα_q/11_ activity with YM-254890 (10 µM), but were unaffected by blocking P/Q-type calcium channels with ω-agatoxin (0.5 µM) (**Fig. 3D, H, I**). Moreover, like the NE-induced bursts of IPSCs, the rhythmic IPSCs generated by hM3D activation in CCK interneurons also underwent transient and complete DSI, which was reversed by the CB1 receptor antagonist AM251 (10 µM) (**Fig. 3J-L**). Collectively, these results confirmed the specificity of the pharmacological manipulation of CCK basket cell-mediated synaptic release and demonstrated that Gq activation in CCK basket cells is both necessary and sufficient to induce NE-mediated inactivating bursts of low-frequency IPSCs.

**Figure 3.**
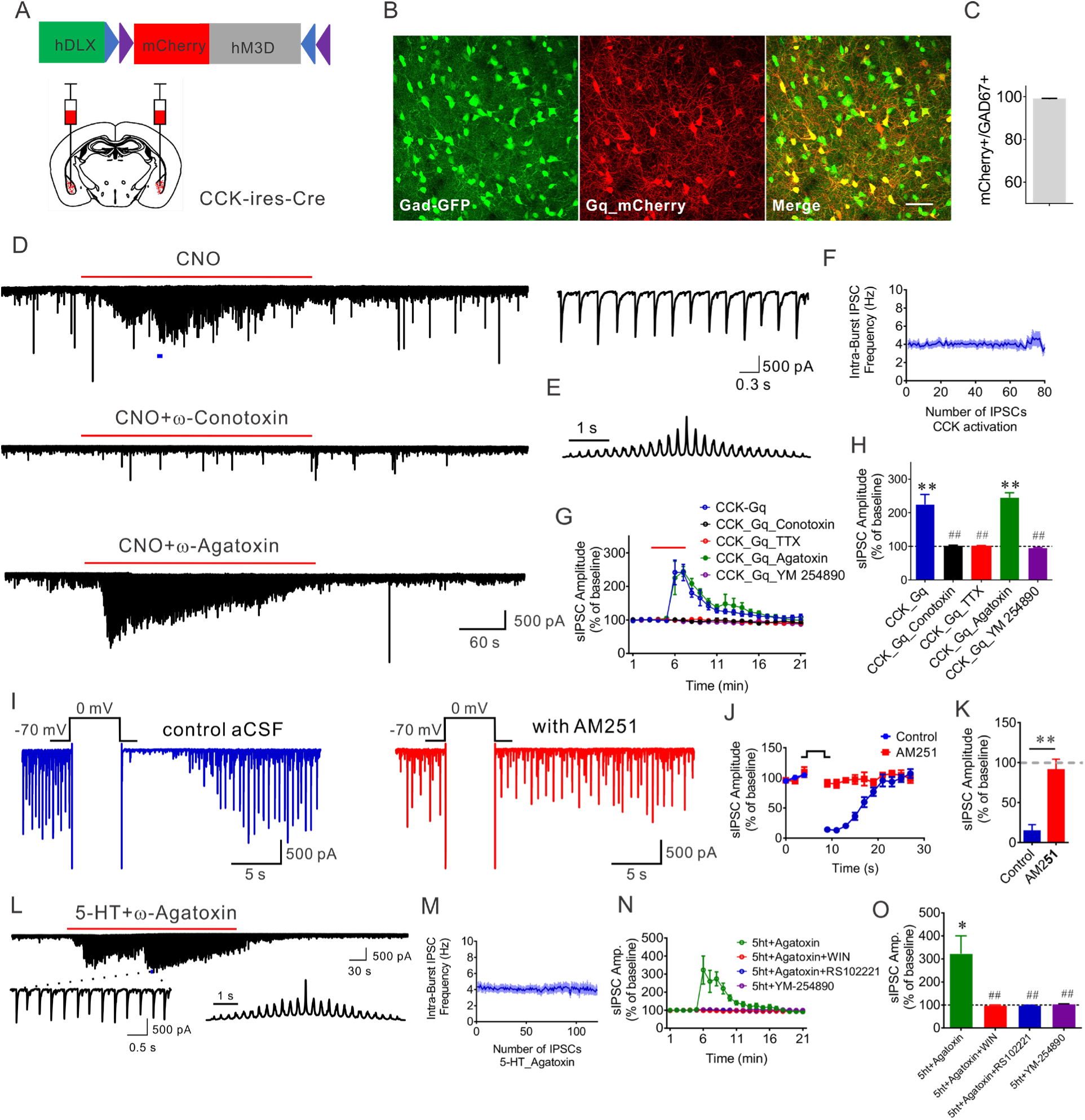
Gq-DREADD activation in BLA CCK interneurons stimulates IPSC bursts in BLA principal neurons. **A**. Schematic diagram showing injection of Cre-dependent hDLX-DIO-hM3D(Gq)-mCherry virus into the BLA of the CCK-ires-Cre mouse. **B.** Representative images of colocalization of hM3D-mCherry and Gad67-GFP in the BLA. **C.** Quantification of hM3D-mCherry and Gad67-GFP co-localization (98.82±0.56%, 1722 cells). **D.** Representative recordings from principal neurons showing that Gq activation in BLA CCK interneurons stimulated bursts of rhythmic, low-frequency IPSCs, which were blocked with the N-type calcium channel blocker ω-conotoxin and unaffected by the P/Q-type calcium channel blocker ω-agatoxin (in different cells). Inset: Expanded trace from the segment in D1 indicated by the bar. **E.** Autocorrelogram showing the rhythmicity of the CNO-induced IPSCs in the recording shown in D1. **F.** Population mean (± SEM) of instantaneous frequency of CNO-induced trains of IPSCs showing constant ∼4-Hz IPSC frequency for the duration of the bursts (n = 13 bursts from 10 cells). **G, H.** Time course (G) and mean change (H) in sIPSC amplitude stimulated by hM3D activation in CCK interneurons. The CNO-induced increase in sIPSC amplitude (10 cells from 5 mice) was blocked by the N-type calcium channel antagonist ω-conotoxin (5 cells from 2 mice), the sodium channel blocker TTX (6 cells from 2 mice), and the selective Gα_q/11_ inhibitor YM-254890 (5 cells from 3 mice), but not by the P/Q-type calcium channel antagonist ω-agatoxin (5 cells from 2 mice). Paired t test, CCK-Gq vs. baseline, p = 0.0022; CCK-Gq-Agatoxin vs. baseline, p = 0.0006, **, p < 0.01; One way ANOVA, F (4, 26) = 10.99, P < 0.0001; Dunnett’s multiple comparisons test, CCK-Gq vs. CCK-Gq-Conotoxin, p = 0.0014; CCK-Gq vs. CCK-Gq-TTX, p = 0.0008; CCK-Gq vs. CCK-Gq-Agatoxin, p = 0.91; CCK-Gq vs CCK-Gq-YM-254890, p = 0.0011, ##, p < 0.01. **I.** Representative recordings of CCK interneuron hM3D-induced IPSCs in a principal neuron showing a depolarization-induced suppression of inhibition (Control) that was reversed by blocking CB1 receptors (AM251), revealing endocannabinoid-dependent DSI. **J, K.** Average time course of (J) and mean change in (K) the amplitude of CNO-induced rhythmic IPSCs in the principal neurons by DSI (9 cells from 4 mice). The DSI is reversed completely by the CB1 receptor antagonist, AM251 (5 cells from 2 mice). Unpaired Student’s t test, control vs. AM251, p < 0.0001, **, p < 0.01. **L.** Representative trace showing that, in the presence of ω-agatoxin (to block PV neuron-mediated bursts), serotonin induced bursts of IPSCs in BLA principal neurons similar to those elicited by NE application and CCK neuron hM3D activation. Insets: Expanded trace and autocorrellogram showing the rhythmicity of the 5-HT-induced IPSCs. **M.** Population mean (± SEM) of instantaneous frequency of 5-HT-induced trains of IPSCs, which shows the constant intra-train IPSC frequency. **N, O.** Time course (N) and mean change (O) in sIPSC amplitude induced by serotonin (5-HT). The 5-HT-induced IPSCs (isolated by ω-agatoxin, in 8 cells from 3 mice) were blocked by: CB1 receptor activation with WIN 55,212-2 (6 cells from 3 mice), the 5-HT2C receptor antagonist RS102221 (6 cells from 3 mice), and the Gq G-protein inhibitor YM-254890 (6 cells from 3 mice). Paired t test, 5HT-Agatoxin vs. baseline p = 0.024; One-way ANOVA, F (3, 24) = 7.836, p = 0.0018; Dunnett’s multiple comparisons test, 5-HT+Agatoxin vs. 5-HT+Agatoxin+WIN, p = 0.0025; 5-HT+Agatoxin vs. 5-HT+Agatoxin+RS102221, p = 0.005; 5-HT+Agatoxin vs. 5-HT+YM-254890, p = 0.006. *, P < 0.05; ##, p < 0.01.

Since both Gq-coupled α1A adrenoreceptor and Gq-DREADD activation in BLA CCK interneurons induced similar bursts of rhythmic, low-frequency IPSCs in the principal neurons, we next asked whether this Gq-dependent bursting mechanism is generalized to other neuromodulators whose cognate Gq-coupled receptors are expressed in CCK interneurons. Serotonin is one of the stress-related neuromodulators that play critical roles in modulating inhibitory neurotransmission in the BLA via Gq-coupled 5-HT receptors (Ji et al., 2017; Jiang et al., 2009; Wang et al., 2021), so we tested for a similar IPSC bursting response to serotonin. Surprisingly, in BLA slices treated with ω-agatoxin (0.5 µM) to block the PV neuron-mediated repetitive IPSC bursting response and with the ionotropic glutamate receptor antagonists DNQX (20 µM) and APV (40 µM) to block excitatory synaptic transmission, bath application of serotonin (5-HT, 100 µM) stimulated multiple inactivating trains of low-frequency rhythmic IPSCs (2.55 to 7.35 Hz, mean = 4.37 Hz) (**Fig. 3M, N**), which were similar to those observed in response to CCK neuron α1A adrenoreceptor and hM3D activation. Blockade of action potentials with TTX (0.5 µM) and CCK interneuron-mediated GABA release with the CB1 receptor agonist WIN 55,212-2 (1 µM) totally abolished the serotonin-induced IPSC trains (**Fig. 3 O, P**), which indicated that the serotonin-induced IPSCs are also generated by activation of presynaptic CCK interneurons. Additionally, pretreatment of slices with selective antagonists for the Gq-coupled 5-HT2C receptor, RS102221 (10 µM), and for the Gα_q_ G-protein inhibitor, YM-254890 (10 µM), also blocked the serotonin-induced increase in IPSCs (**Fig. 3 O, P**). Thus, multiple different neuromodulators converge onto a Gq signaling mechanism in CCK neurons to generate a common rhythmic, synchronized inhibitory output to regulate BLA principal neuron activity. We next tested for an effect of the synchronous Gq signaling-induced bursting inhibitory synaptic input to BLA principal neurons on BLA output.

### Gq signaling in BLA CCK basket cells promotes BLA network state switches *in vivo*

Given the influence of interneurons on the coordination of local network activity in the BLA, we postulated that Gq activation in CCK interneurons may be involved in reconfiguring BLA population-level activity *in vivo*. To test this, we recorded local field potentials in the BLA *in vivo* in CCK-ires-Cre mice expressing Gq-DREADD in BLA CCK interneurons via unilateral BLA injections of virus (AAVdj-hDLX-DIO-hM3D-mCherry) (**Fig. 4A**). Consistent with the observation that Gq activation in the CCK interneurons generated rhythmic IPSCs at ∼4 Hz in BLA principal cells, we observed that activation of the Gq-DREADD in BLA CCK interneurons with CNO infusion into the BLA (1 mM, 0.5 μl) significantly increased oscillations in the low (2-5 Hz) and high theta range (6-12 Hz) (**Fig. 4B-D**). Interestingly, CNO administration also decreased the BLA oscillation power in the slow (40-70 Hz) and fast gamma range (80-120 Hz) (**Fig. 4B, F, G**), which could be due to Gq activation of other types of CCK-expressing interneurons in the CCK-ires-Cre mouse line including the PV interneurons, which generate phasic bursts of IPSCs (**Suppl. Fig. 2A**) and decrease slow and fast gamma in response to Gq activation (Fu et al., 2022).

**Figure 4.**
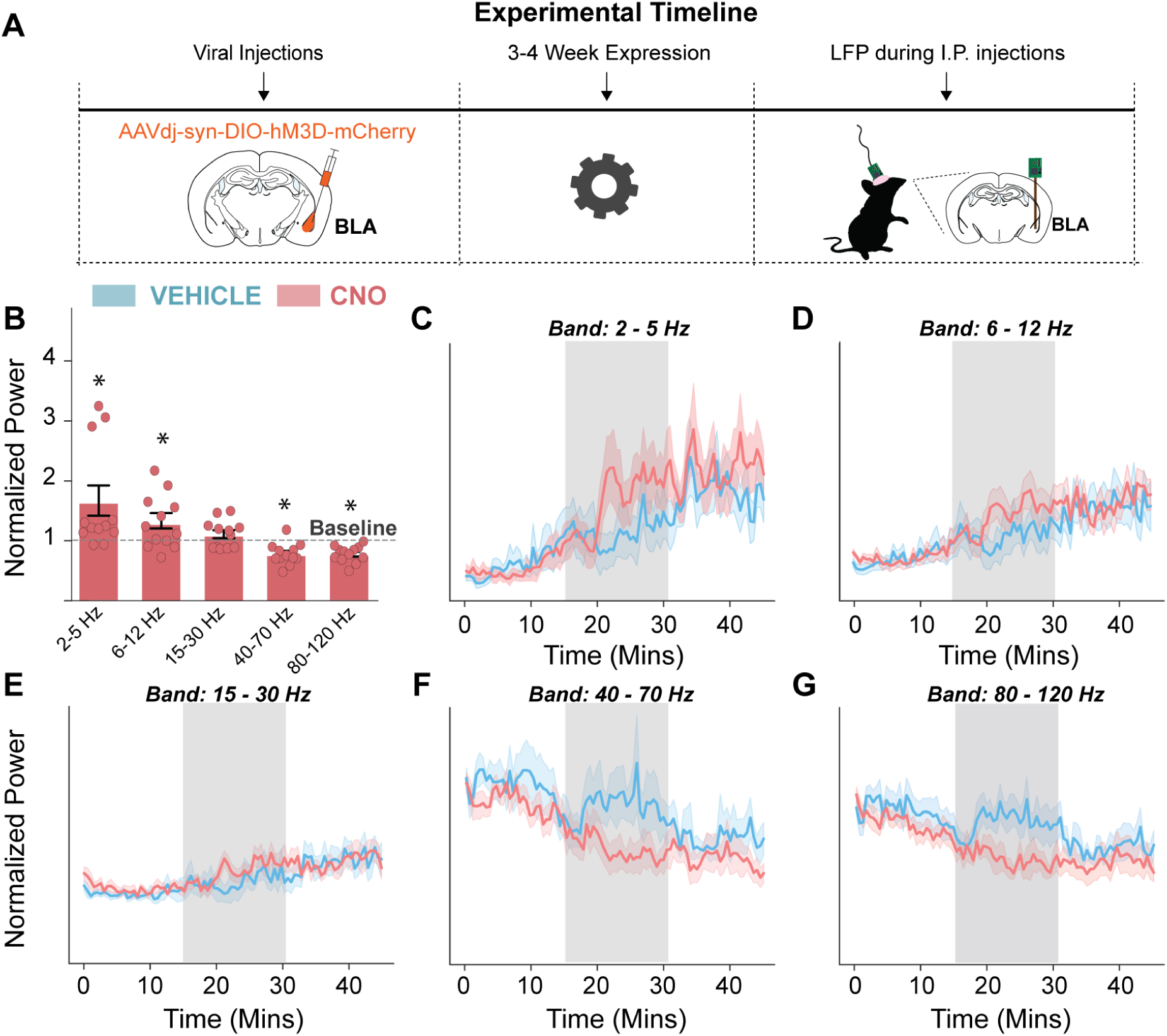
Chemogenetic activation of BLA CCK interneurons modulates oscillations in the BLA. **A.** Schematic of the experimental timeline. **B.** Mean (±SEM) normalized power of oscillations in the BLA across frequencies. Individual values were represented as the power in response to CNO infusion compared to vehicle (n = 12 mice). Two-tailed, one-sample *t* test, 2-5 Hz, p = 0.0312, 6-12 Hz, p = 0.0498, 15-20 Hz, p = 0.1927, 40-70 Hz, p = 0.0008, 80-120 Hz, p < 0.0001, *, p < 0.05. **B-F.** Power across time for 2-5 Hz (C), 6-12 Hz (D), 15-30 Hz (E), 40–70 Hz (F), and 80–120 Hz (G) normalized to baseline for both vehicle (blue) and CNO (red) infusions. Blue/red lines = average power over time, blue/red-shaded areas = SEM.

### Activation of Gq signaling in CCK interneurons suppresses conditioned fear learning

Stressful stimuli trigger a significant increase in NE release in the BLA, which plays a key role in the modulation of fear memory formation (Galvez et al., 1996; McGaugh, 2004; Quirarte et al., 1998). To investigate the role in fear memory formation of α1A adrenergic receptor and Gq-DREADD activation of synchronized rhythmic inhibitory signals from CCK interneurons to principal neurons in the BLA, we used a site- and cell type-specific α1A adrenoreceptor rescue and Gq-DREADD expression strategy in the global adra1AKO mouse. Thus, we tested the behavioral effects of re-expression of α1A adrenoreceptors or Gq-DREADD activation specifically in BLA CCK interneurons with Cre-dependent hDLX intersectional AAV virus delivered bilaterally in the BLA of adra1AKO mice in a standard auditory-cued fear conditioning paradigm (**Fig. 5A**).

**Figure 5.**
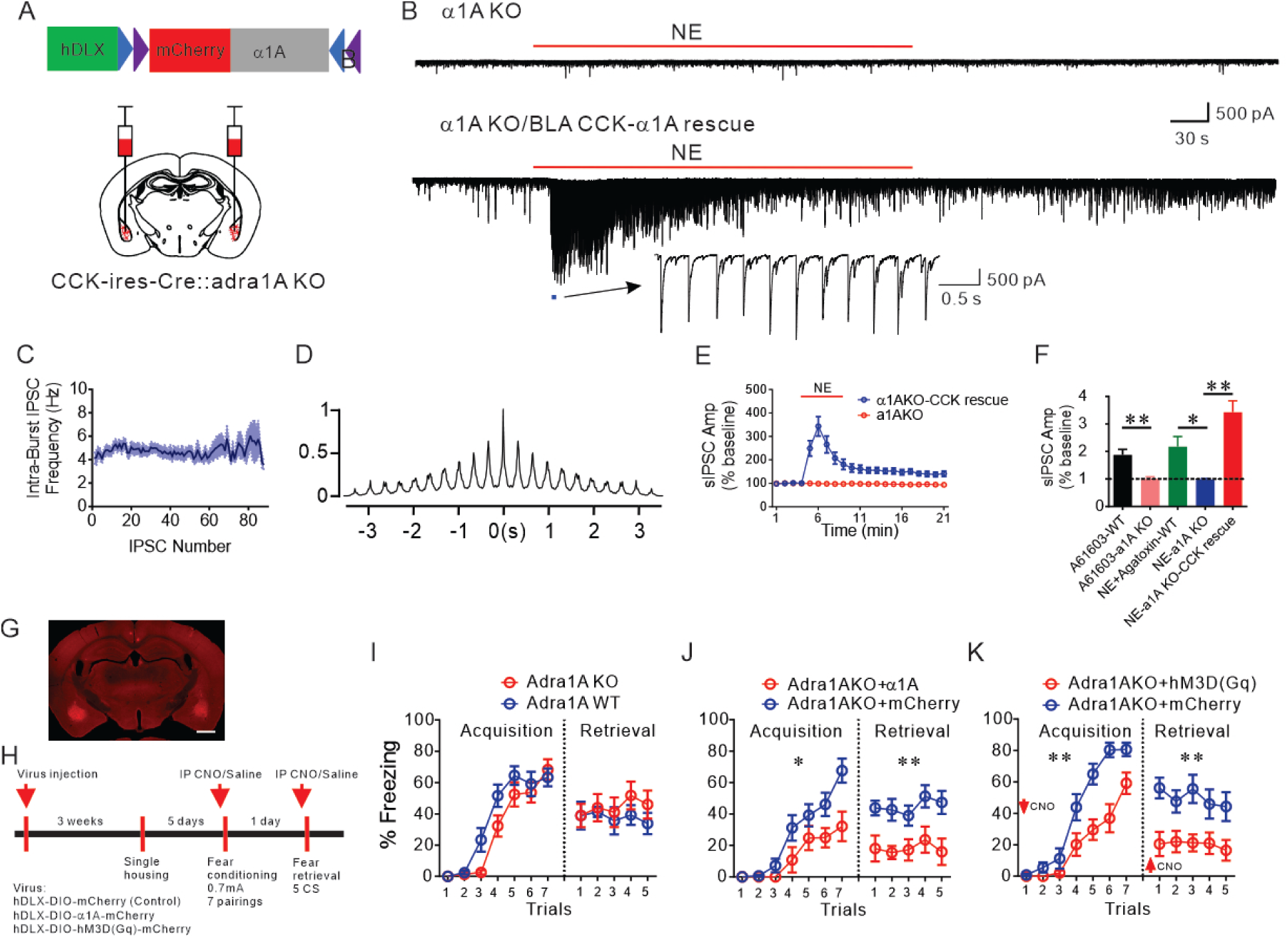
α1A adrenoreceptor restoration and Gq-DREADD activation in BLA CCK interneurons rescues rhythmic IPSC trains and suppresses fear memory. **A.** Diagram showing injection of Cre-dependent hDLX-α1A virus into the BLA of CCK-ires-Cre::adra1A KO mice. **B.** Representative traces showing the lack of response to NE in a BLA principal neuron in a slice from a global adra1A KO mouse (Top trace) and restoration of the NE-induced trains of rhythmic IPSCs following re-expression of the α1A adrenoreceptors in the CCK interneurons (Bottom trace). **C.** Population mean (± SEM) of the instantaneous frequency of the rescued CCK-mediated IPSCs (8 bursts from 8 cells). **D.** Autocorrelogram showing the rhythmicity of the restored CCK-mediated IPSCs. **E.** Time course of the NE effect on sIPSC amplitude in adra1A KO mice with and without virally-transduced re-expression of α1A adrenoreceptors in CCK interneurons. **F.** Mean change in sIPSC amplitude in BLA principal neurons in response to NE or the α1A agonist A61603 in slices from wild type (WT) and adra1A KO mice. (NE+Agatoxin_WT: 10 cells from 4 mice; NE_adra1A KO: 9 cells from 4 mice; A61603_WT: 10 cells from 4 mice; A61603_adra1A KO: 7 cells from 3 mice; NE_adra1A KO_CCK rescue: n = 9 cells from 3 mice) (Unpaired t test, A61603_WT vs. A61603_adra1A KO: p = 0.0032, ** p < 0.01; One-Way ANOVA, F (2, 25) = 13.85, p < 0.0001, Dunnett’s multiple comparisons test, NE+Agatoxin_WT vs. NE_adra1A KO: p = 0.0268, NE_adra1A KO vs. NE_adra1A KO_CCK rescue: p < 0.0001, * p < 0.05, ** p < 0.01). **G.** A representative image showing bilateral injection of the conditional AAV-DLX virus in the BLA to label the CCK interneurons. Scale bar, 1 mm. **H.** Experimental timeline for virus injection and fear conditioning. CCK-ires-Cre::adra1A KO mice were injected with conditional AAVs expressing Cre-dependent α1A receptors or Gq-DREADD to rescue Gq-coupled receptor signaling in BLA CCK interneurons. Control CCK-ires-Cre::adra1A KO mice were injected with conditional AAVs expressing only mCherry. For Gq-DREADD rescue experiments, CNO (5 mg/kg) was administrated I.P. 30 min prior to commencement of the fear acquisition and again the next day 30 min prior to commencement of the fear retrieval (red arrows) in both Gq-DREADD and mCherry groups. **I.** Global adra1A deletion did not change fear memory acquisition (Repeated measures Two-Way ANOVA, F (1, 20) = 2.49, p = 0.13) and recall compared to wildtype littermate controls. (Repeated measures Two-Way ANOVA, F (1, 20) = 0.60, p = 0.45). **J.** Rescue of α1A adrenoreceptors selectively in BLA CCK interneurons significantly attenuated fear acquisition (Repeated measures Two-Way ANOVA, F (1, 21) = 6.63, p = 0.018 compared to control virus-injected CCK-ires-Cre::adra1A KO group), and the retrieval of the fear memory (Repeated measures Two-Way ANOVA, F (1, 21) = 20.73, p = 0.0002). * p < 0.05, ** p < 0.01 compared to control virus-injected CCK-ires-Cre::adra1A KO group. **K.** Gq signaling in BLA CCK interneurons in adra1A KO mice with excitatory DREADD decreased fear memory acquisition (F (1,20) = 17.69, p = 0.0004) and fear memory retrieval (Repeated measures Two-Way ANOVA, F (1,20) = 16.26, p = 0.0007). ** p < 0.01 compared to control virus injected in CCK-ires-Cre::adra1A KO.

Prior to testing for the role of CCK α1A adrenoreceptors in fear conditioning, we first validated our receptor rescue strategy by testing with whole-cell recordings in *ex vivo* brain slices from the adra1A global knockout mouse for the effect of α1A adrenoreceptor knockout and re-expression in BLA CCK interneurons on the NE-induced CCK neuron-mediated rhythmic IPSC bursts. In slices from the adra1AKO mouse with excitatory transmission blocked with DNQX (20µM) and APV (40µM), NE (100 µM) and the selective α1A adrenoreceptor agonist A61603 (2 µM) failed to elicit an increase in sIPSCs (**Fig. 5B, E, F**), confirming the specific α1A adrenoreceptor dependence of the NE-induced IPSC burst generation. After two weeks following injection of the Cre-dependent, α1A-expressing hDLX virus (AAVdj-hDLX-DIO-α1A-mCherry) bilaterally into the BLA of CCK-ires-Cre::adra1A KO mice, NE activation of α1A adrenoreceptors re-expressed in CCK interneurons in the adra1AKO mouse rescued the bursts of low-frequency rhythmic IPSCs in the BLA principal neurons (n=9 cells from 4 mice) (**Fig. 5B-F**). In DNQX (20 µM) and APV (40 µM) to block excitatory synaptic transmission, NE (100 µM) predominantly induced inactivating ‘bursts’ of rhythmic, low-frequency IPSCs in the BLA principal neurons with an instantaneous intra-burst frequency of 4-5 Hz (range: 2.82 to 6.73 Hz, mean: 4.6 Hz) and a duration of 80 s (range: 28 to 170.81 s, mean: 79.98 s). In a subset of recorded neurons (2 out of 9 cells), we also observed repetitive bursts of IPSCs with accelerating intra-burst frequency that were likely to be mediated by Gq activation in PV interneurons, in addition to the CCK interneuron-mediated rhythmic IPSCs (**supplement Fig. 2B**), which is similar to hM3D activation of CCK interneurons in the CCK-ires-Cre mice. Overall, these results suggested that rescue of α1A adrenoreceptor signaling in CCK-ires-Cre::adra1A KO restored rhythmic IPSCs mediated by NE activation of CCK interneurons.

After validating the effectiveness of the virus-based rescue of NE-induced CCK-type inhibitory synaptic inputs to BLA principal cells, we next investigated the role of α1A adrenergic engagement of CCK basket cells in modulating fear memory formation. Three weeks after re-expressing α1A adrenoreceptors with bilateral injection of AAVdj-hDlx-DIO-α1A-mCherry into the BLA of CCK-ires-Cre::adra1A KO mice (**Fig. 5A, G**), the mice underwent a standard auditory-cued fear conditioning paradigm (**Fig. 5H**). Surprisingly, mice with global knockout of α1A adrenoreceptors did not show significant differences in fear memory acquisition or retrieval compared with their wildtype littermate controls (adra1A WT vs. adra1A KO: fear acquisition, p=0.13; fear retrieval, p = 0.45) (**Fig. 5I**). In contrast, selective rescue of α1A adrenergic signaling specifically in BLA CCK interneurons resulted in impaired fear acquisition and retrieval, as evidenced by reduced freezing behavior during both fear conditioning training (p = 0.018) and subsequent presentation of the auditory tone alone 24 h later (p = 0.0002) (**Fig. 5J**).

We next tested the effects of Gq-DREADD activation in BLA CCK interneurons on fear learning in the adra1AKO mice. Three weeks after bilateral injection of AAVdj-hDLX-DIO-hM3D(Gq)-mCherry or mCherry control virus into the BLA of CCK-ires-Cre::adra1A KO, CNO (5 mg/kg) was administered I.P. 30 min prior to beginning the fear acquisition paradigm and again 30 min prior to beginning the fear retrieval test the next day. Like the effects of α1A adrenoreceptor re-expression in the adra1AKO mouse, Gq activation in CCK interneurons also inhibited fear acquisition (p = 0.0004) and retrieval (p = 0.0007) (**Fig. 5K**). This suggested, therefore, that the induction of bursts of rhythmic IPSCs and altered BLA network states by α1A adrenoreceptor or Gq-DREADD activation in CCK interneurons in the BLA suppresses associative fear memory formation and retrieval.

## DISCUSSION

Although cannabinoid modulation of inhibitory synaptic transmission mediated by CB1-expressing CCK basket cells has been proposed to regulate the stress facilitation of fear memory formation (Atsak et al., 2015; Campolongo et al., 2009; Morena et al., 2016), direct investigation of the role of CCK interneuron activity stimulated by stress-related neuromodulators in learned fear has been largely impeded by the lack of cell type-specific genetic tools targeting the CCK interneurons. Here, we demonstrate that CB1-expressing CCK interneurons are activated by Gq-coupled receptors to generate a rhythmic-patterned inhibitory input to the BLA principal neurons that modulates BLA network states and fear memory formation. With an interneuron-specific, Cre-dependent hDLX virus delivered to CCK-Cre-expressing mice, we found that the selective re-expression of α1A adrenoreceptors or expression of Gq-DREADD in the BLA CCK interneurons of global adra1A knockouts and the resulting rescue of rhythmic perisomatic IPSCs in BLA principal neurons and altered BLA network states suppresses the acquisition and expression of conditioned fear memory.

Despite the previously documented role of NE in the facilitation of inhibitory synaptic transmission in the BLA (Braga et al., 2004; Miyajima et al., 2010), knowledge of the identities of the presynaptic interneurons and the patterns of inhibitory synaptic activation by NE has been limited. According to our double-dissociation experiments of NE-induced IPSCs with calcium channel blockers and a CB1 receptor agonist, the inactivating bursts of low-frequency rhythmic IPSCs in BLA principal neurons are mediated by NE activation of presynaptic CCK interneurons. We showed in a previous study that NE also stimulates a distinct repetitive bursting pattern of high-frequency inhibitory postsynaptic currents in the BLA principal neurons that is mediated by activation of a second subtype of presynaptic inhibitory interneuron, the PV basket cells (Fu et al., 2022). Consistent with the interpretation here that NE-induced low-frequency, rhythmic IPSC bursts are mediated by CCK interneurons, the IPSC bursts underwent complete suppression by CB1 receptor activation and DSI, which are dependent on the CB1-expressing inhibitory synaptic terminals of CCK basket cells (Freund et al., 2003; Malhotra et al., 2025). Moreover, like PV interneurons (Fu et al., 2022), the NE-induced CCK neuron-generated IPSC bursts were mediated by activation of α1A adrenoreceptors. We found that most of the CCK interneurons in the BLA (∼75%) express α1A adrenoreceptors and that restoration of the CCK interneuron α1A adrenoreceptors in the adra1A KO mouse rescued the NE-induced IPSC trains. Taken together, these data show that α1A noradrenergic activation in CCK interneurons generates synchronized bursts of rhythmic low-frequency IPSCs in BLA principal neurons.

The NE stimulation of CCK interneuron-mediated IPSC bursts was α1 adrenoreceptor-dependent and inhibited by the selective Gα_q/11_ inhibitor YM-254890, suggesting that it is mediated by activation of the Gq G-protein signaling pathway. A surprising finding of our study was that selective activation of Gq signaling in the CCK interneurons with the hM3D DREADD generated similar bursts of rhythmic IPSCs that were sensitive to N-type calcium channel blockade and cannabinoid receptor activation. We found that the CB1-sensitive bursts of rhythmic IPSCs were also stimulated by serotonin activation of Gq-coupled 5-HT2C receptors. Therefore, the bursting response of the CCK neurons was not specifically dependent on the neurotransmitter or the neurotransmitter receptor *per se*, but rather on the G-protein signaling pathway engaged. We found previously that the bursting activation of PV basket cells is also G protein dependent and agnostic to receptor subtype (Fu et al., 2022). Interestingly, a recent study in the BLA (Bratsch-Prince et al., 2024) and multiple studies in the CA1 subfield of the hippocampus (Alger et al., 2014; Nagode et al., 2011; Nagode et al., 2014) have found that activation of Gq-coupled M1/3 cholinergic receptors also induce similar ∼4-Hz, CB1-sensitive rhythmic IPSCs from CCK interneurons. It is worth noting that, like NE inputs, both serotonin and acetylcholine circuits are also activated during arousal and regulate BLA output and emotional behaviors (Crimmins et al., 2023; Crouse et al., 2020; Yu et al., 2022). Thus, the generation of bursts of rhythmic IPSCs in the principal neurons may serve as the functional output of Gq activation in the CCK interneurons by multiple neuromodulators transmitting emotional salience. Since the BLA is a cortical-like brain structure and similar rhythmic inhibitory synaptic activity has been reported in the hippocampus and neocortex, it is likely that this stereotyped response to Gq signaling in CCK neurons generalizes to other cortical structures.

Selective genetic access to the CCK interneurons is hindered by the fact that low levels of CCK or its preprohormone are expressed in principal neurons (Taniguchi et al., 2011). To circumvent this problem, we used an intersectional strategy combining Cre-driven recombination in CCK-expressing neurons and GABA neuron specificity of viral expression with the hDLX promoter (Dimidschstein et al., 2016; Liu et al., 2020) to target the CCK interneurons. After re-expressing the α1A adrenoreceptors in the CCK interneurons bilaterally in the BLA in the CCK-ires-Cre::adra1A KO mouse, and following *ex vivo* confirmation of the rescue of the Gq-induced IPSC bursting in the principal neurons, we found that chemogenetic activation of CCK interneurons enhanced theta oscillations in the BLA in vivo, similar to a previous report (Bratsch-Prince et al., 2024), and suppressed the acquisition and expression of associative fear learning. These data suggest that CCK interneuron-mediated enhancement of theta rhythms in the BLA is associated with suppression of fear memory formation. This inhibitory effect of CCK interneuron activation on conditioned fear is consistent with a prior study that showed that 20 Hz optogenetic stimulation of CCK interneurons during conditioning facilitated subsequent fear extinction (Rovira-Esteban et al., 2019). Relative to a recent study showing that whole-brain Gq activation in CCK interneurons promotes fear learning (Whissell et al., 2019), our data provide brain region specificity and suggest that CCK interneuron activation in the BLA may produce opposite effects to those in other areas of the brain, consistent with the opposing regulatory effects of the amygdala and prefrontal cortex/hippocampus on stress circuits (Herman et al., 2016; Ulrich-Lai & Herman, 2009).

In the neocortex, hippocampus, and BLA, synchronized rhythmic perisomatic inhibition is critically involved in the generation of neural circuit oscillations. It well established that the rhythmic IPSCs induced by muscarinic receptor activation in CCK interneurons drives theta oscillations in CA1 (Alger et al., 2014; Nagode et al., 2011; Reich et al., 2005), and recent evidence suggests a similar phenomenon in the BLA (Bratsch-Prince et al., 2024). As would be predicted from the synchronized rhythmic IPSCs at ∼4 Hz, we found that α1A noradrenergic activation in CCK interneurons increased the power of theta oscillation in the BLA. Emerging evidence has shown that oscillation at theta frequency in the BLA entrained from the prefrontal cortex is essential for the expression of learned fear (Bocchio et al., 2017; Karalis et al., 2016). Interestingly, we found that the intrinsic CCK neuron Gq-mediated rhythmic bursts at theta frequency in the BLA suppressed fear acquisition and expression. One possible explanation for the opposite effects of extrinsic and intrinsic theta oscillation on fear expression is phase interference, by which the intrinsically driven CCK basket cell-mediated rhythmic IPSCs might disrupt the timing of the top-down PFC-driven BLA theta oscillations, leading to an inhibition of fear expression. Further, the extrinsic engagement of theta oscillations in the BLA that is not time locked to the stimulus may prevent the establishment of the association between CS and US necessary for fear learning.

CCK-positive GABAergic interneurons are heterogeneous, including both large-somata, CB1-positive basket cells and small-somata, CB1-negative cells that co-express vasoactive intestinal polypeptide (VIP) (Jasnow et al., 2009; Katona et al., 2001; Mascagni & McDonald, 2003; Vogel et al., 2016). As a result, CCK interneurons targeted with injection of the DLX virus in CCK-ires-Cre mice should include both types of cells. Gq activation in the entire population of BLA CCK-expressing interneurons generated IPSCs in the principal cells that underwent strong DSI, with nearly 100% suppression, which suggests that the principal neurons we recorded received few, if any, inputs from CCK-positive, CB1-negative VIP interneurons. This is consistent with BLA VIP interneuron projections preferentially to other interneurons and not directly to the principal neurons (Krabbe et al., 2019; Rhomberg et al., 2018). Functionally, the VIP interneurons have been shown to provide disinhibitory gating to the BLA principal cells and activation of these cells enhanced fear learning (Krabbe et al., 2019). Thus, whereas activation of the VIP interneurons alone facilitated associative fear learning, our findings demonstrated that activation of the entire CCK interneuron population, including both the CCK basket cells and the CCK/VIP cells, inhibits fear memory acquisition. In addition to the CCK population heterogeneity, it has also been reported that intersectional targeting of the CCK interneurons with CCK-ires-Cre mice also labels some PV-positive interneurons (Rovira-Esteban et al., 2019). Since restoration of α1A adrenoreceptor activation in BLA PV interneurons in the ara1AKO mouse facilitates fear expression (Fu et al., 2022), the inhibitory effect of Gq activation in CCK-ires-Cre mice on fear learning suggests that any effect of Gq activation of PV interneurons is dwarfed by activation of the CB1-positive CCK basket cells. The Gq activation in PV interneurons may contribute to the suppression of gamma oscillations we observed following Gq-DREADD activation of the BLA CCK-expressing interneurons, since we found that Gq activation of BLA PV neurons in the ara1AKO mouse suppressed gamma (Fu et al., 2022). Taken together, these data indicate that CCK basket cells in the BLA enhance theta and suppress gamma oscillations, and restrain fear memory formation by generating bursts of rhythmic IPSCs in the principal neurons in response to NE. Whether the effect on fear memory retrieval is caused by inhibition of fear acquisition, of fear expression, or both will require further studies to separate the effects of Gq activation on fear acquisition and retrieval.

Compared with the accelerating and repetitive bursts of IPSCs generated by Gq activation in PV interneurons (**Suppl Fig. 1** and Fu et al., 2022), the CCK interneuron-mediated IPSCs driven by Gq activation exhibited a stable frequency and were clustered in a single, inactivating train. Thus, the same ligands (NE, CNO, and serotonin) acting on the same types of receptors (hM3D, α1A adrenoreceptors, and 5-HT2A/5-HT2C serotonergic receptors, respectively) produced distinct patterns of activation in CCK and PV interneurons. Notably, Gq-coupled receptor activation in these two BLA basket cell subpopulations had opposite effects on fear learning: Gq activation in PV interneurons facilitated fear retrieval, whereas Gq activation in CCK interneurons suppressed both fear acquisition and retrieval. These cell type-specific responses may result from differences in electrophysiological properties and intracellular signaling targets between the two cell types (Armstrong & Soltesz, 2012). The contrasting effects on BLA oscillations and behavioral outputs, despite similar perisomatic projections, likely arise from their distinct bursting activities and the resulting differential regulation of principal cell firing patterns. Since neuromodulators often act via volume transmission to influence populations of neurons, this dichotomy in output patterns from basket cells in response to Gq activation by the same ligands may provide a mechanism for neuromodulators to diversify their regulation of BLA neural circuits. Determining how neuromodulators achieve specificity in regulating principal neurons, despite converging on common signaling mechanisms, remains an important question for future study.

NE α1A adrenergic signaling in CCK interneurons led to inhibition of conditioned fear acquisition and expression, which is opposite the effect in PV neurons, in which α1A adrenoreceptor activation has a facilitatory effect on fear learning (Fu & Tasker, 2024; Fu et al., 2022). This opposing effect of noradrenergic modulation of PV and CCK neurons may explain why there was no overall effect of global adra1A knockout on fear learning. Overall, the diversified responses of PV and CCK interneurons to the same neuromodulator presents a possible mechanism by which stress can fine tune associative fear learning by balancing the contribution of different patterns of perisomatic inhibition. A question for future study is under what contexts are NE actions on PV interneurons and/or CCK interneurons engaged and what is their corresponding impact on network and behavioral states.

In conclusion, we present a cellular mechanism for controlling brain oscillations and fear memory formation through a rhythmic inhibitory synaptic input to BLA principal neurons by Gq-coupled receptor activation in CCK interneurons. The result of this CCK neuron inhibitory input differs dramatically from the effect of phasic input from PV neurons generated by similar Gq-coupled receptor activation. Thus, during emotional arousal, Gq activation by diverse neuromodulatory signals may titrate the contribution of different patterns of perisomatic inhibitory inputs to the BLA principal cells to achieve the appropriate fear memory regulation, and disruption of this signaling balance could contribute to neuropsychiatric disorders.

---

**Supplemental Figure 1. Norepinephrine elicits two patterns of IPSC bursts that are mediated by activation of different subtypes of presynaptic inhibitory interneuron. A1.** The two patterns of NE-elicited IPSC bursts: an initial, single inactivating burst of low-frequency IPSCs followed by a second series of repetitive bursts of high-frequency IPSCs that outlasted the NE application. **A2.** The repetitive high-frequency IPSC bursts, but not the initial low-frequency IPSC bursts, were blocked by the P/Q calcium channel blocker ω-agatoxin. **A3.** The initial low-frequency burst was blocked by the CB1 receptor agonist WIN 55,212-2 (WIN). **B.** Time histogram of IPSC amplitudes elicited by NE, normalized to baseline IPSC amplitudes. The long-lasting response was blocked by the P/Q calcium channel blocker ω-agatoxin and the initial response was blocked by CB1 receptor activation with WIN (NE, n=16 cells; NE+Aga, n=10 cells; NE+Aga+WIN, n=8 cells). **C.** Population mean and SEM of intra-burst IPSC frequency of the repetitive, high-frequency IPSC bursts as a function of the relative time position within the bursts (38 bursts from 16 cells). IPSC frequencies accelerated to a relatively high frequency (33.05 +/-2.08 Hz) at the beginning of the bursts before gradually slowing until termination of the bursts.

**Supplemental Figure 2. Gq activation in CCK interneurons with CCK-ires-Cre mice by hM3D or restored α1A receptors in adra1A KO mice induced different types of IPSC bursts in a subset of recorded BLA principal neurons. A, B.** Representative recordings of IPSCs in principal neurons during CNO activation of hM3D in a CCK-ires-Cre mouse (A) or NE activation of restored a1A adrenoreceptors in a CCK-ires-Cre::adra1AKO mouse (B) showing two types of IPSC bursts. In addition to the rhythmic IPSC bursts at the beginning (expanded panel on the left), repetitive bursts of IPSCs with accelerating frequency (expanded panel on the right) were also observed in a subset of neurons (∼20%).

