## Supplemental figures 1 and 2. for "Noradrenergic neuromodulation of cholecystokinin interneurons in the basolateral amygdala alters rhythmic activity and restrains fear memory"

**SUPPLEMENTAL FIGURE 1**

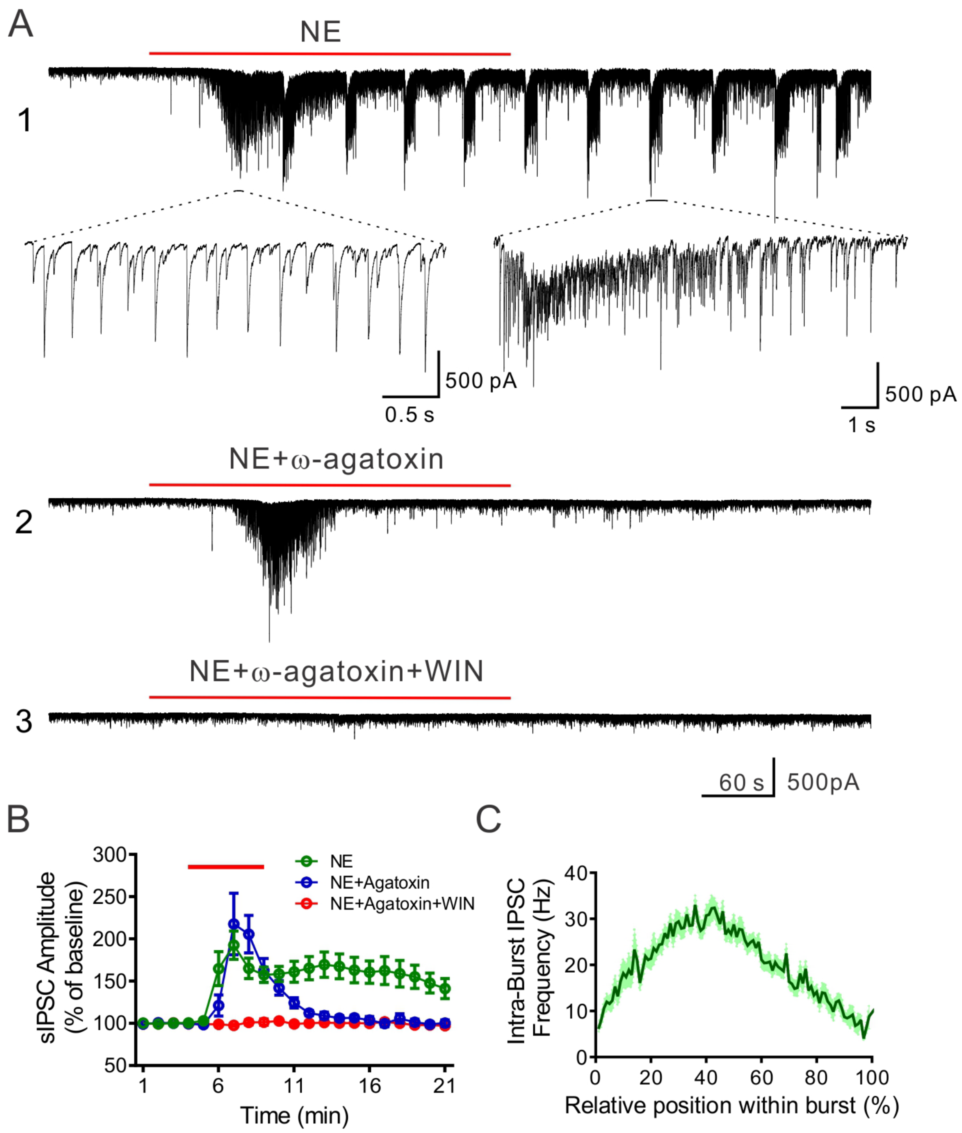


**Supplemental Figure 1. Norepinephrine elicits two patterns of IPSC bursts that are mediated by activation of different subtypes of presynaptic inhibitory interneuron. A1.** The two patterns of NE-elicited IPSC bursts: an initial, single inactivating burst of low-frequency IPSCs followed by a second series of repetitive bursts of high-frequency IPSCs that outlasted the NE application. **A2.** The repetitive high-frequency IPSC bursts, but not the initial low-frequency IPSC bursts, were blocked by the P/Q calcium channel blocker ω-agatoxin. **A3.** The initial low-frequency burst was blocked by the CB1 receptor agonist WIN 55,212-2 (WIN). **B.** Time histogram of IPSC amplitudes elicited by NE, normalized to baseline IPSC amplitudes. The long-lasting response was blocked by the P/Q calcium channel blocker ω-agatoxin and the initial response was blocked by CB1 receptor activation with WIN (NE, n=16 cells; NE+Aga, n=10 cells; NE+Aga+WIN, n=8 cells). **C.** Population mean and SEM of intra-burst IPSC frequency of the repetitive, high-frequency IPSC bursts as a function of the relative time position within the bursts (38 bursts from 16 cells). IPSC frequencies accelerated to a relatively high frequency (33.05 +/- 2.08 Hz) at the beginning of the bursts before gradually slowing until termination of the bursts.

**SUPPLEMENTAL FIGURE 2**

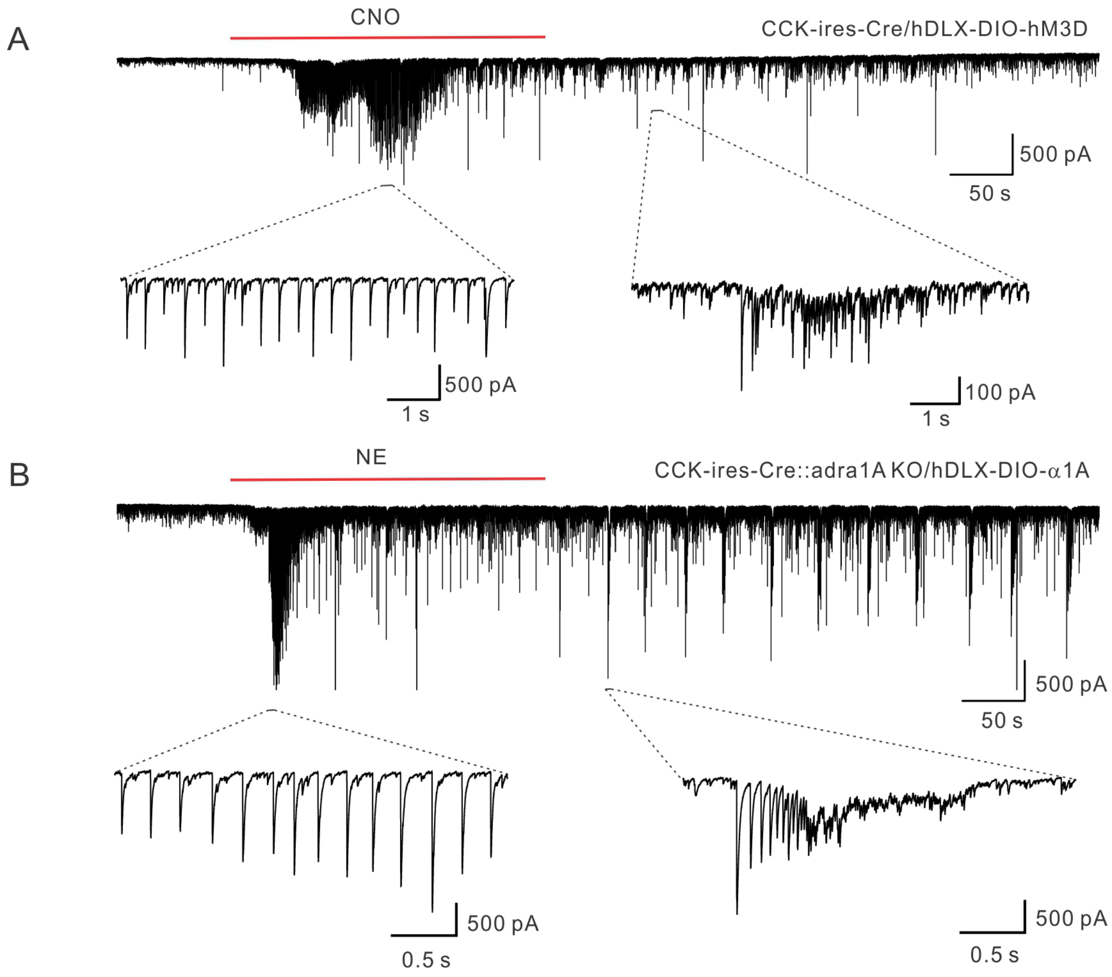


**Supplemental Figure 2. Gq activation in CCK interneurons with CCK-ires-Cre mice by hM3D or restored α1A receptors in adra1A KO mice induced different types of IPSC bursts in a subset of recorded BLA principal neurons. A, B.** Representative recordings of IPSCs in principal neurons during CNO activation of hM3D in a CCK-ires-Cre mouse (**A**) or NE activation of restored a1A adrenoreceptors in a CCK-ires-Cre::adra1AKO mouse (**B**) showing two types of IPSC bursts. In addition to the rhythmic IPSC bursts at the beginning (expanded panel on the left), repetitive bursts of IPSCs with accelerating frequency (expanded panel on the right) were also observed in a subset of neurons (~20%).
